# Extracellular vesicles from saxitoxin-producing strains of the cyanobacterium *Raphidiopsis raciborskii* and investigation of their potential allelopathic effect

**DOI:** 10.1101/2025.07.02.662845

**Authors:** Cezar Paiva do Nascimento, Mauro Cesar Palmeira Vilar, Paulo Mascarello Bisch, Ana Beatriz Furlanetto Pacheco

## Abstract

*Raphidiopsis raciborskii* is a filamentous cyanobacterium with global distribution in tropical and temperate climates. Toxic strains isolated in Latin America produce saxitoxins (STXs), which are neurotoxins that block voltage-gated ion channels in cells. This cyanobacterium can exert allelopathic effects on other phytoplankton species in response to resource competition through the secretion of substances that impair the target-cell physiology. Extracellular vesicles (EVs) constitute a secretion system that can transport biologically active molecules to distant targets in a protected, concentrated and targeted way. However, the role of EVs in cyanobacterial biology is poorly understood. We aimed to characterize *R. raciborskii* EVs and test whether this extracellular fraction or other exudate fractions contribute to its allelopathic effect. Exudates from two STX-producing *R. raciborskii* strains (T3 and CY-10) were subjected to filtration and divided into vesicular fractions (VF, >100 kDa, with EVs), which were concentrated by ultracentrifugation, and dissolved fractions (DF, <100 kDa, no EVs). Characterization of the VF through nanoparticle tracking analysis and transmission electron microscopy revealed spheric structures with diameters of 100-200 nm. Ratios of the number of EVs/cell were estimated as 21.54 ± 1.33 for T3 and 8.79 ± 0.38 for CY-10. The STX content of exudate fractions was determined by ELISA. Total extracellular STX concentrations (STX + neosaxitoxin (neoSTX)) were 170.64 (±15.43) pg.mL^-1^ for T3 and 913.84 (±226.87) pg.mL^-1^ for CY-10 and these values were similar to those of the DF (<100 kDa fraction). In the VF, STX concentrations were approximately 10³–10⁴ times lower. The estimation of toxin quotas (per biovolume) resulted in values 20-40 times lower in EVs (0.27±0,09 ng.mm^-3^ for T3 and 1.81±0.49 ng.mm^-3^ for CY-10) than in cells (13.05 ±2.16 ng.mm^-3^ for T3 and 33.65 ±7.31 ng.mm^-3^ for CY-10). A shift in the predominant analog between compartments was observed, with neoSTX prevailing in cells and STX in EVs. The CY-10 strain exhibited higher toxin concentrations than T3 in the VF, DF, total extracellular fraction and cellular fraction. The allelopathic effect of *R. raciborskii* exudate fractions was tested on the green algae *Monoraphidium capricornutum*. The total extracellular fraction and the DF equally inhibited *M. capricornutum* growth and photosynthesis, but the VF had no effect. Thus, the effect can be attributed to dissolved allelopathic compounds <100 kDa. These results open new opportunities to further investigate the role of EVs in the ecophysiology of *R. raciborskii*.

**Highlights:**

- Extracellular vesicles (EVs) from two toxic strains of *R. raciborskii* were characterized;
- Saxitoxins can be released within EVs;
- Extracellular saxitoxins content includes dissolved and vesicular fractions;
- While cyanobacterial EVs had no allelopathic effect the dissolved extracellular fraction (<100 kDa) of *R. raciborskii* did.

## Introduction

*Raphidiopsis raciborskii* (Order Nostocales) is a freshwater cyanobacterial species with a wide geographic distribution and invasive potential. Its adaptive success can be related to diverse aspects such as production of gas vesicles, which allow migration in the water column and better positioning in the photic zone, nitrogen fixation, resistance in adverse conditions (differentiation of akinetes), efficient uptake and storage of phosphorus and allelopathic potential (Antunes et al., 2015; Burford et al., 2016). Toxic strains of *R. raciborskii* isolated from South America produce the neurotoxin saxitoxin while those isolated from Asia and Oceania produce the cytotoxin cylindrospermopsin (Lagos et al., 1999; Antunes et al., 2015, Burford et al., 2016). Saxitoxins (STXs) are a group of guanidine alkaloids consisting of about 60 variants that block ion channels (Na^+^ and Ca^2+^) and modulate potassium channels (K^+^) in eukaryotic cells. The analogues of STXs are classified into three main groups, based on the number of SO_3_H groups present in their structures: di-sulfated (C-toxins), mono-sulfated (gonyautoxins, GTXs) or non-sulfated groups, saxitoxins (STXs) (Cusick & Sayler, 2013; Wiese et al., 2010). STX is an intracellular toxin, but a smaller portion can be exported, presumably through the putative multidrug and toxic compound extrusion family transporters SxtF and SxtM (Soto-Liebe et al., 2013).

The transport of cytoplasmic components to the external medium or to other cells is an important aspect of bacterial adaptation to the environment. There are several classes of bacterial secretion systems composed of multi protein complexes that cross a single phospholipid membrane, two membranes, or even three membranes, involving the bacterial cell itself and a target cell (Green & Mecsas, 2016). The secretion of selected molecules enables detoxification, adhesion, motility and competition, among other functions. There is yet another type of transport system called the type zero secretion system, which is based on extracellular vesicles (Guerrero-Mandujano et al., 2017). Extracellular vesicles (EVs) are released by cells from all domains of life. In bacteria, these structures are delimited by a lipid membrane and have a diameter of 50 to 250 nm, but vary in their biogenesis, structure and content (Toyofuku et al., 2023). In Gram-negative bacteria, blebbing of the outer membrane can generate membrane vesicles carrying periplasmic content. Alternatively, vesicles with both membranes can form when gaps in the peptidoglycan layer allow the protrusion of the inner membrane, in this case they also carry cytosolic content. Besides, in both Gram-negative and Gram-positive bacteria, membrane vesicles can originate from cell lysis carrying material from all cell compartments (Toyofuku et al., 2023). EVs are an efficient mechanism to deliver bioactive molecules to target cells, preventing their degradation and reaching greater distances than other bacterial secretion systems. Multiple functions have been described for bacterial EVs, such as relief from envelope stress, phage defense, nutrient acquisition, DNA transfer and cellular communication, among others (Guerrero-Mandujano et al., 2017; Lima et al., 2020; Nagakubo et al., 2020).

For cyanobacteria, little is known about EVs compared to other bacteria (Lima et al., 2020). EVs have been mainly detected in unicellular morphotypes representative from marine systems, such as *Synechococcus* (Biller et al., 2014, 2023; Xu et al., 2013; Yin et al., 2019), *Prochlorococcus* (Biller et al., 2014, 2023), *Synechocystis* (Oliveira et al., 2016; Pardo et al., 2015) and *Cyanothece* (Mota et al., 2020). Regarding filamentous morphotypes, production of EVs was observed in the freshwater genera *Anabaena* (Oliveira et al., 2015), *Raphidiopsis* (Zarantonello et al., 2018), *Leptolyngbya* (Usui et al., 2022) and in the marine species *Jaaginema litorale* (Brito et al., 2017). However, most studies focused on EV characterization, while their functional role remains elusive. The production of EVs by a STX-producing *R. raciborskii* strain (CYRF) has been previously reported (Zarantonello et al., 2018). In addition to the characterization of EV morphology by electron microscopy, the authors suggested that vesiculation would be an adaptive response to stress, increasing upon UV radiation exposure and interaction with other cyanobacterial species. Overall, proposed functions for bacterial EVs in aquatic environments involve defense from phages, nutrient fluxes and nucleic acid transfer (Biller et al., 2014; 2017; 2023).

Cyanobacterial metabolites can exert allelopathic effects in phytoplankton components (Antunes et al., 2015; Śliwińska-Wilczewska et al., 2018, 2021). Allelopathy can be defined as any direct or indirect (negative or positive) effect of chemicals produced and secreted by an organism on another organism (Rice, 1985). In aquatic environments, inhibitory chemical interactions represent an important aspect of phytoplankton ecology (Legrand et al., 2003). The primary effects are inhibition of growth and photosynthetic activity on microalgal targets (Leão et al., 2009b; Śliwińska-Wilczewska et al., 2018). These effects have been tested in culture conditions, showing that <u>cyanobacterial exudates</u> contain bioactive compounds that inhibit phytoplankton organisms (Leão et al., 2009b; Rzymski et al., 2014; Chia et al., 2021). Among other secondary metabolites, cyanotoxins can act as allelochemicals (Leão et al., 2009b; Teneva, et al., 2023). *R. raciborskii* exudates, in particular, showed allelopathic inhibition of the chlorophytes and other cyanobacterial species (Figueredo et al., 2007; Leão et al., 2009a; Antunes et al., 2012; Rzymski et al., 2014). Therefore, allelopathy is considered a relevant ecological interaction influencing phytoplankton dynamics and algal bloom establishment (Śliwińska-Wilczewska et al., 2021). Although bacterial EVs have been recognized as algicidal to microalgae (Li et al., 2024), their involvement in cyanobacterial allelopathic interactions has not been tested, similarly, the presence of toxins in cyanobacterial EVs has not been investigated.

Here, we aimed to characterize *R. raciborskii* EVs and investigate their role in toxin release and allelopathy. Since STXs are actively exported to the surrounding and may be involved on cyanobacterial allelopathy, we hypothesized that i) EVs could participate in the secretion of this toxin and ii) play a role on *R. raciborskii* exudate allelopathic effect on other phytoplankton organisms.

## Materials and Methods

### Microorganisms and culturing conditions

Two saxitoxin-producing *Raphidiopsis raciborskii* strains were used: CY-10 (isolated from the São Sebastião do Alto reservoir, RJ, Brazil, 21°57’26" S 42°08’05" W in 2018) and T3 (isolated from the Billings reservoir, SP, Brazil, 23°47’13"S, 46°35’2"W in 1996). The profile of saxitoxin variants produced by T3 has been described previously (Lagos et al., 1999; Soto-Liebe et al., 2010). For CY-10, the saxitoxin profile was described in the present study. The cyanobacterial strains were obtained from the collection of the Laboratory of Ecophysiology and Toxicology of Cyanobacteria (Federal University of Rio de Janeiro, RJ, Brazil). The chlorophyte *Monoraphidium capricornutum* (formerly *Selenastrum capricornutum*) was kindly provided by the Ecotoxicology Laboratory of the Mineral Technology Center (CETEM-RJ). Cultures were maintained in ASM-1 medium (Gorham et al., 1964), under non-axenic conditions, at a constant temperature of 24 ± 0.5°C, with a photon flux of 30 µmol m⁻² s⁻¹ for cyanobacteria and 80 µmol m⁻² s⁻¹ for the chlorophyte and a 12-hour photoperiod.

For the experimental cultures, for each *R. raciborskii* strain, cultures were initiated with an inoculum of 20 mm³.L⁻¹, with four biological replicates of 1.8 L in 2L Erlenmeyers, at 24 ± 0.5°C, 30 µmol m⁻²× s⁻¹, 12-hour photoperiod. Cultures were maintained until cells reached the exponential phase. Cultures of *Monoraphidium capricornutum* were established in ASM-1 and initiated with a cell density corresponding to at 40 µg.L⁻¹ of chlorophyll-α, maintained in 150 mL Erlenmeyer flasks filled with 70 mL of ASM-1 culture medium. Each condition included 4 four replicates.

### Growth curve and biovolume determination

The growth of *R. raciborskii* in culture was monitored by collecting samples every two days over a 14-day period. Samples were preserved in 1% acetic Lugol solution. Trichome counting was carried out in a Fuchs-Rosenthal chamber using an Olympus BX51 optical microscope and for image acquisition the Cell^B software package was used (Olympus). The resulting trichome density was converted to biovolume (mm³.L⁻¹) based on the average trichome volume (µm³), as described by Hillebrand et al. (1999) and Sun & Liu (2003). The average trichome volume was calculated from the measurement of the length and diameter of 150 trichomes for T3 and 190 trichomes for CY-10.

Specific growth rate (*µ*) was calculated during the exponential growth phase using the following equation (1) (Fogg & Thake, 1987)

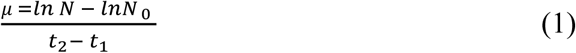

Where *N* = final biovolume; *N_0_* = initial biovolume; *t*_2_ = final time; and *t*_1_ = initial time.

Cyanobacterial growth rates were calculated based on biovolume measurements while for the chlorophyte growth rates were calculated based on chlorophyll concentrations.

### Cyanobacterial culture fractioning and extracellular vesicles obtaining

Extracellular fractions and cellular fractions were obtained from *R. raciborskii* cultures at the exponential phase, as determined from growth curves (10-12 days). At this point, the cultures presented a biovolume of 121 mm³×L⁻¹ and 108 mm³×L⁻¹ for CY-10 and T3, respectively. The vesicle extraction was conducted separately for each replicate culture as described (Biller et al., 2022) (a scheme of the procedure is shown in Supplementary material S1).

Prior to EV purification the total culture volume was centrifuged at 4,000 ×g for 15 minutes at 10 °C to separate intracellular (cell pellet) and extracellular (exudate) fractions. The supernatant (hereafter exudate) was collected, and the cell pellet was stored at -20 °C. Using a Kitasato connected to a vacuum pump, the exudate was filtered through a 0.45 µm pore membrane and then through a 0.22 µm pore membrane using 45 mm filters (Merck Millipore). The filtrate corresponded to the total extracellular (referred to as EF), which was also stored at -20 °C.

To concentrate the vesicles, the filtrate from the 0.22 µm pore membrane was submitted to tangential filtration using the Vivaflow® 200 device (Sartorius) with a 100 kDa membrane. The volume that passed through the membrane represented the dissolved fraction (referred to as DF), with particles <100 kDa and without vesicles, which was stored at -20 °C. The concentrated volume (containing vesicles and other particles > 100 kDa) was subjected to two ultracentrifugation steps at 120,000 ×g (Beckman TL100 centrifuge) at 5 °C for 1 hour. The supernatant was discarded, and the pellet was resuspended in 300 µL of phosphate buffer (PBS). This volume was then filtered through a 0.22 µm pore membrane (Merck Millipore), and this fraction was designated as the vesicular fraction (VF). The samples were stored in PBS at -20 °C.

### Nanoparticle Tracking Analysis

Nanoparticle Tracking Analysis (NTA) was used to calculate the concentration of EVs and to evaluate their hydrodynamic diameter. The NanoSight equipment (NS300) (Hardware: embedded laser: at 532 nm; camera: sCMOS) was set according to the manufacturer’s software manual (User Manual, MAN0541-02-EN, 2018). Five 60-s videos were captured under the following conditions: temperature of 25°C; syringe speed of 25 µl/s, camera level 13, screen gain 1.0. After capture, the videos were analysed by the in-build NanoSight Software (NTA 3.4 Build 3.4.4) with a detection threshold of 7. The VF preparations were diluted in PBS at different ratios, ranging from 1:50 to 1:250, until an appropriate reading pattern was obtained.

Based on the average diameter value and particle concentration, the total vesicle biovolume was calculated by assuming the volume of a sphere and multiplying by the total number of vesicles.

### Transmission Electron Microscopy

To obtain microscopy images of EVs, 400 mesh copper grids were coated with a formvar film. The procedures were performed as described by Biller et al., 2022. Briefly, a volume of 50 μL of sample was placed on the grid and, after 30 seconds, part of the liquid was removed from the edge using a piece of filter paper. Subsequently, 50 μL of uranyl acetate (saturated aqueous solution) was added. After 30 seconds, the excess of uranyl acetate was removed with filter paper and the grid was left to dry at room temperature (25 °C). Samples were analysed using a HITACHI HT7800 electronic microscope at the National Center for Structural Biology and Bioimaging (CENABIO), Federal University of Rio de Janeiro.

### Analysis of Extracellular Saxitoxins by Enzyme-Linked Immunosorbent Assay (ELISA)

STX and neoSTX were quantified by ELISA in the extracellular (EF), dissolved (DF), and vesicular (VF) fractions. Prior to the analysis, VF underwent lysis through three freeze-thaw cycles followed by ultrasound sonication (50 Hz for 15 minutes). STX quantitation was achieved using competitive ELISA kits (96 wells) from Beacon Analytical Systems, according to the manufacturer’s instructions. Results were expressed in µg STX_eq_.L⁻¹, with a quantification limit of 0.02 µg STX_eq_.L⁻¹. These values were used to calculate the toxin quota considering cell or EV biovolumes (ng.mm^-3^).

### Analysis of Saxitoxins by High Performance Liquid Chromatography – Refractive Fluorescence Detection

Saxitoxin extraction from *R. raciborskii* cell biomass was performed as described by Vilar et al. (2021). Toxins were analyzed using high-performance liquid chromatography (Shimadzu, Class VP) with a fluorescence detector (HPLC-RFD). A C18 reverse-phase column and a 20 µL injector were employed. Chromatographic analyses followed Oshima (1995) post-column derivatization method with a mobile phase of 2 mM sodium 1-heptanesulfonate in 30 mM ammonium phosphate and 5% acetonitrile for non-sulfated saxitoxins and 2 mM sodium 1-heptanesulfonate in 10 mM ammonium phosphate for mono-sulfated saxitoxins (gonyautoxins) at a flow rate of 0.8 mL.min⁻¹. STXs were detected at 330 nm excitation and 400 nm emission. The STX standards used were: saxitoxin (STX), decarbamoyl neosaxitoxin (dcSTX), neosaxitoxin (neoSTX), and gonyautoxins (GTXs 1-5), obtained from the National Research Council, Institute of Marine Biosciences (Canada). The total STX concentration was expressed in ng STX.mL⁻¹. The limit of quantification of the method was 0.55 ng, 0.004 ng, 1.01 ng, 3.43 ng, 0.242 ng, 2.08 ng, 0.5 ng, 0.239 and 1.30 ng for neoSTX, dcSTX, STX, GTX 1-5, respectively. Also, STX content per cellular biovolume was estimated as a saxitoxins quota per biomass (ng.mm^-3^).

Saxitoxin concentrations from *R. raciborskii* exudates were also determined by this method to use this measurement as a basis to test its allelopathic effect.

### Allelopathic effect of Raphidiopsis raciborskii exudate fractions or saxitoxins on Monoraphidium capricornutum

Cultures of the algae *Monoraphidium capricornutum* were initiated at 1.2×10^5^ cells.mL^-1^ corresponding to ∼40 µg.L⁻¹ of chlorophyll-α, (as described in “Microorganisms and culturing conditions”) and employed to test the allelopathic effect of the different *R. raciborskii* exudate fractions: VF, EF and DF **(Figure 1a)**.

**Figure 1:**
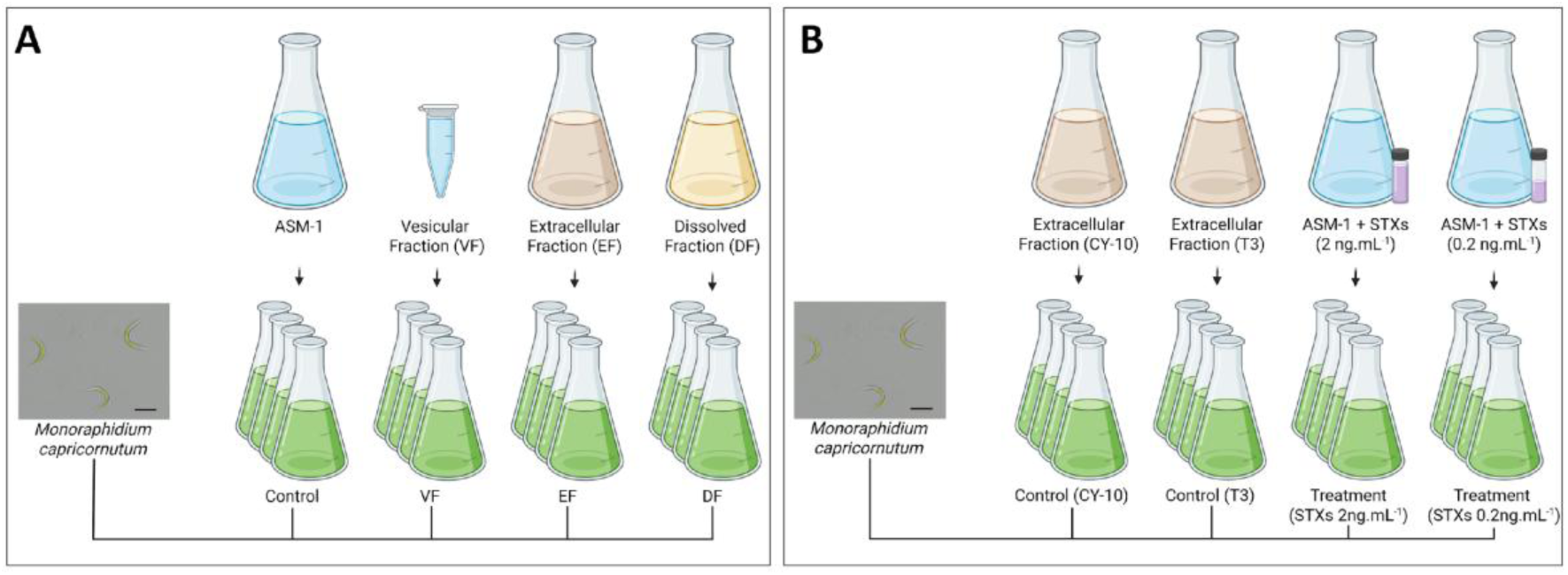
(A) Scheme of the experimental set-up to test the allelopathic effect of *Raphidiopsis raciborskii* exudate fractions on *Monoraphidium capricornutum* (scale bar 10 µm). (B) Scheme of the experimental set-up to test *M. capricornutum* response to purified saxitoxins at the concentrations similar to those detected in the exudates of the different studied *R. raciborskii* strains. Each condition included 4 replicates.

To test whether purified cyanobacterial EVs contribute to the exudate allelopathic effect on the chlorophyte, an aliquot of concentrated VF (in PBS) was added to *M. capricornutum* cultures. The final EV concentration at which the chlorophyte was exposed was 20-fold higher than that measured in the original cyanobacterial cultures. In the control condition, 1 mL of PBS (resulting in 1.43% PBS in ASM-1) was added to *M. capricornutum* cultures.

To test the effect of the extracellular and dissolved fractions (EF and DF, with or without EVs, respectively), these fractions were first added with the ASM-1 salts (NaNO₃, MgSO₄, MgCl₂, CaCl₂, KH₂PO₄, and Na₂HPO₄) to to keep macronutrients at levels similar to those found in standard ASM-1 culture medium and avoid any side-effect due to inorganic nutrient limitation. *M. capricornutum* cultures were established in these modified ASM-1.

Moreover, assuming that STXs in the exudates of *R. raciborskii* play a role on its allelopathy, we also tested the effect of purified STX analogues, similar to those produced by the tested cyanobacterial strains, on *M. capricornutum* **(Figure 1b)**. Purified STXs analogues (obtained from the National Research Council, Institute of Marine Biosciences, Canada) were added to the cultures at mean concentrations corresponding to those measured in the cyanobacterial exudates: 2 ng.mL⁻¹ for *R. raciborskii* CY-10 (comprising 80% neoSTX, 10% dcSTX, and 10% STX) and 0.2 ng.mL⁻¹ for *R. raciborskii* T3 (50% neoSTX and 50% STX). In the STX assays, extracellular fractions (EF) from *R. raciborskii*, served as a positive control for the allelopathic effect. EF was first added with the ASM-1 salts to keep macronutrients at levels similar to those of the standard culture medium. *M. capricornutum* cultures were established in these modified ASM-1.

Each condition included 4 replicates. Samples from the cultures were taken at 0, 3, and 9 days to analyze photosynthetic parameters and at 0, 1, 3, 6, and 9 days to determine chlorophyll-α concentration.

### Chlorophyll-α concentration and photosynthetic parameters

A PHYTO-PAM fluorometer (Heinz Walz GmbH, Germany) equipped with a PHYTO-EDF detection unit for measuring fluorescence was used to evaluate photosynthetic activity. Fluorescence data, obtained after applying saturation pulses (36 µmol photons m⁻² s⁻¹), were converted into chlorophyll-α concentration (µg.L⁻¹) and photosynthetic yield (relative *Fv’/Fm’*).

Light curves were generated by exposing dark-acclimated culture samples (F₀) to light intensities ranging from 16 to 764 µmol photons m⁻²s⁻¹, with intervals of 10 seconds between pulses. From these curves, the maximum electron transport rate (ETR_max_), light saturation (*Ik*) and light-harvesting efficiency (α) were calculated as functions of irradiance. In addition, the effective quantum yield of photosynthetic energy conversion in PSII, defined as φm = [(*F*_m_ − *F*)/*F*_m_], was determined, where *F* is the fluorescence of the dark-adapted sample and *F*_m_ represents the maximum dark-adapted fluorescence.

### Statistical analysis

All datasets were tested for normality and variance homogeneity and once assumed the parametric premises, the following tests were performed: Student’s *t*-test, one-way and RM two-way ANOVA. To account for multiple comparisons and control Type I error, the Bonferroni *post hoc* correction was applied. The significance level was set at α = 0.05. All analyses were performed using GraphPad Prism 8.0.1.

## Results

### Obtaining and characterization of EVs from R. raciborskii strains

Extracellular vesicles (EVs) were obtained from cultures of *R. raciborskii* strains at the exponential growth phase with growth rates corresponding to 0.209 (± 0.02) d^-1^ for T3 and 0.202 (± 0.02) d^-1^ for CY-10, and no differences between the two strains **(Supplementary Figure 2)**. Biovolume concentration corresponded to 108±7.13 mm³.L⁻¹ for T3 and 121±5.42 mm³.L⁻¹ for CY-10.

Transmission electron microscopy (TEM) confirmed the presence of EVs in the preparations and revealed spheric structures with varying apparent diameters around 100 nm **(Figure 2, Supplementary Figure 3)**. Nanoparticle tracking analysis (NTA) evidenced a similar size distribution of the particles for both examined strains, with diameters ranging from 80 to 350 nm **(Figure 2)**. The median diameter values corresponded to 212 nm for T3 and 197 nm for CY-10.

**Figure 2:**
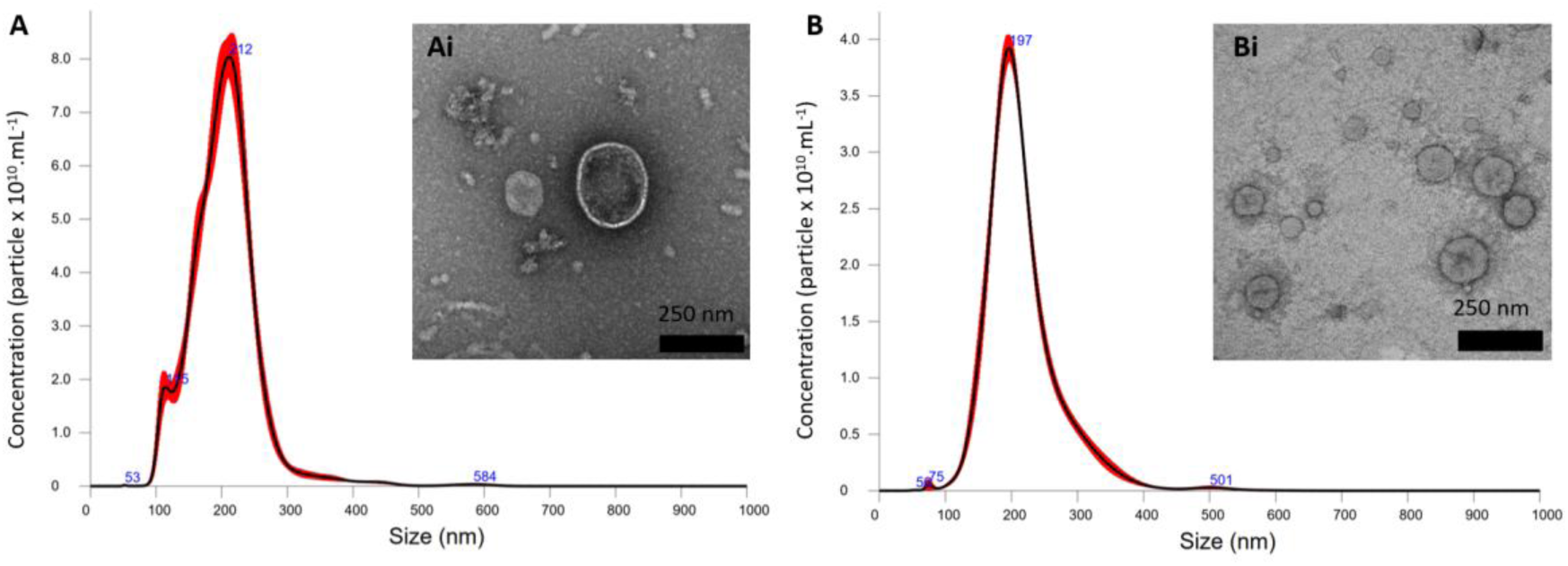
Detection of extracellular vesicles in fractions obtained from cultures of *R. raciborskii* (A) T3 and (B) CY-10 strains. Nanoparticle Tracking Analysis (NTA) provided the size distribution (nm) and concentration (particle x 10^10^.mL^-1^) of particles in EV preparations. Transmission electron microscopy (TEM) images (Ai and Bi) illustrate the typical morphology of the extracellular vesicles.

Considering the biovolume reached by the cyanobacterial strains in the cultures from which EVs were obtained, an estimation of the number of EV per cell was obtained, resulting in 21.54 ± 1.33 for T3 and 8.79 ± 0.38 for CY-10, with a significant difference between the strains **(Supplementary Figure 4A)**.

### Saxitoxin content in R. raciborskii cells and exudate fractions

The *R. raciborskii* strains exhibited unique saxitoxin profiles with different total concentrations and proportions of analogues. In both strains neoSTX was the major analogue (97% for T3 and 83% for CY-10). T3 also produced STX, and CY-10 produced STX and the mono-sulfated GTX-1 and GTX-2 **(Figure 3A)**. The toxin concentration in the intracellular fraction was approximately twice as high in CY-10 as in T3, as determined by volumetric concentrations and by cellular toxin quota per biovolume (ng.mm⁻³) (Student’s *t*-test, *p* < 0.01; **Figure 3A and B)**.

**Figure 3:**
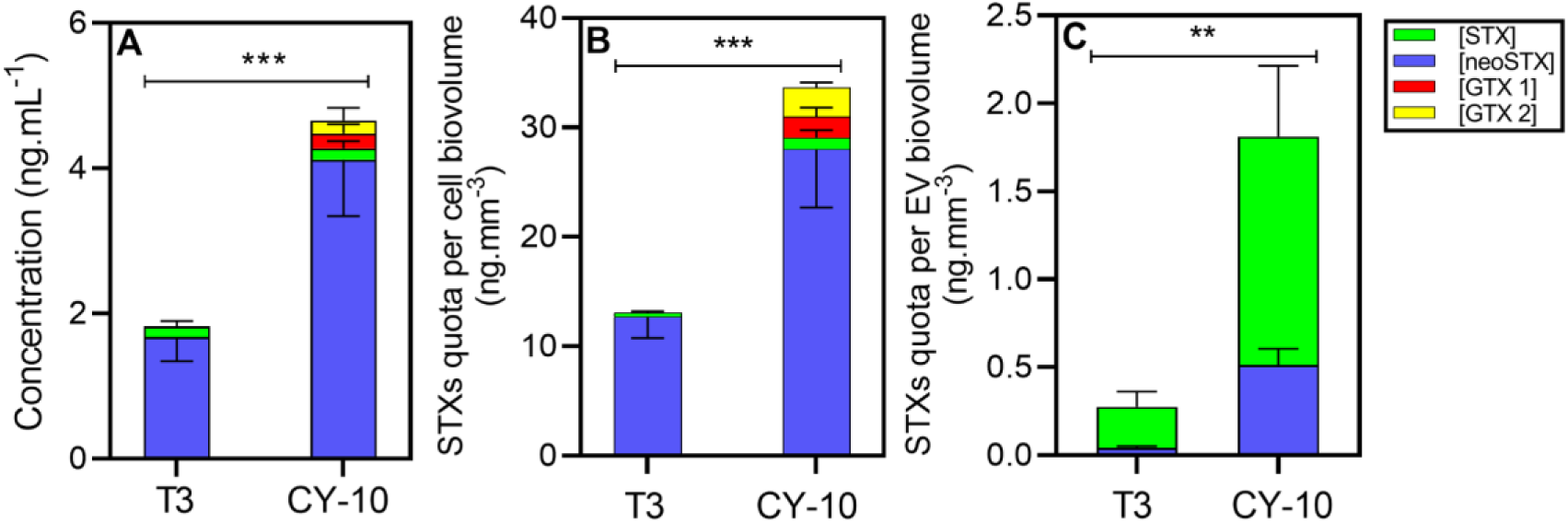
Saxitoxin content of cellular and vesicle fractions of *R. raciborskii* strains. (A) Total intracellular STX concentration produced by *R. raciborskii* T3 and CY-10 strains, measured by HPLC-RFD. (B) STXs quota per cell biovolume. (C) STXs quota per vesicle biovolume. neoSTX: neosaxitoxin; STX: saxitoxin; GTX 1 and 2: gonyautoxins 1 and 2. (**, *** significant differences according to Student’s t-test, *p* < 0.01, *p* < 0.001, respectively).

In the extracellular fractions (EF), STX concentrations were 5-10 times lower than in the cellular fractions **(Table 1)**. The subfraction obtained from the exudate following 100 kDa membrane filtration, named dissolved fraction (DF, <100 kDa fraction), contained STX concentrations similar to those of EFs (Bonferroni’s test; *p* > 0.05) indicating that these toxins are mostly dissolved. The CY-10 strain exhibited 5-fold higher extracellular toxin concentrations than the T3 strain.

**Table 1:** Concentrations of saxitoxin (STX) and neosaxitoxin (neoSTX) measured by ELISA in the exudate fractions of T3 and CY-10 strains. **EF:** total extracellular fraction, **DF:** dissolved fraction and **VF:** vesicular fraction (Mean ± SD). Comparisons among the three extracellular fractions were performed for each strain and different letters indicate significant differences (One-way ANOVA with Bonferroni’s test, *p* < 0.05; Mean ± SD)

| Strain | Fraction | STX (pg.mL<sup>-1</sup>) | neoSTX (pg.mL<sup>-1</sup>) |
| --- | --- | --- | --- |
| T3 | EF | 140.39 (± 10.31)<sup>a</sup> | 30.25 (± 5.12)<sup>a</sup> |
| | DF | 129.31 (± 19.23)<sup>a</sup> | 22.00 (± 4.00)<sup>b</sup> |
| | VF | 0.059 (± 0.02)<sup>b</sup> | 0.01 (± 0.002)<sup>c</sup> |
| CY-10 | EF | 498.17 (± 64.80)<sup>a</sup> | 415.67 (± 162.07)<sup>a</sup> |
| | DF | 465.99 (± 28.09)<sup>a</sup> | 342.33 (± 69.61)<sup>a</sup> |
| | VF | 0.15 (± 0.09)<sup>b</sup> | 0.04 (± 0.01)<sup>b</sup> |

In the present study, cyanotoxins (saxitoxins) were first detected in the vesicular fraction (VF) composing the *R. raciborskii* exudate. In VF, STX concentrations were much lower than those in the other fractions (approximately 10³–10⁴ times lower), but still within the detection and quantification limit of ELISA. Considering that vesicles present a very small volume in relation to the cell biovolume (0.1-0.2%, **Supplementary figure 4B)**, the toxin concentration was converted as quota per vesicle biovolume for comparison purposes. Toxin quotas in vesicles were approximately 20 times lower than those in cells **(Figure 3C)**. A shift in the predominant analog between compartments was observed, with neoSTX prevailing in cells and STX in vesicles. The CY-10 strain exhibited higher toxin content in the VF than the T3 strain (2.7 times more), as observed for total exudate, dissolved and cellular fraction.

### Allelopathic effect of exudate fractions from R. raciborskii cultures

Growth and photosynthesis of the green microalgae *Monoraphidium capricornutum* were monitored toward exposure (9 days) to total extracellular fractions (EF), dissolved fractions (DF), and vesicular fractions (VF) obtained from each STXs-producing *R. raciborskii* strain.

Total (EF) and dissolved (DF <100 kDa) fractions of the two toxic *R. raciborskii* strains significantly reduced *M. capricornutum* growth (as assessed by chlorophyll-α concentration) **(Figures 4A and 5A).** A decrease in the final algal biomass after 9 days was also observed in the cultures exposed to *R. raciborskii* exudate fractions. Similar effects were observed for both fractions **(Table S1)**. For *M. capricornutum* cultures exposed to *R. raciborskii* T3 exudate fractions, the growth rate decreased to 0.24 ± 0.007 d^-1^ and 0.23 ± 0.009 d^-1^ for EF and DF treatments, respectively, compared to 0.42 ± 0.01 d^-1^ in the control (Bonferroni’s test; *p* < 0.05). Regarding the effect of *R. raciborskii* CY-10 exudate fractions, an even more remarkable growth reduction was observed: 0.13 ± 0.05 d^-1^ in the presence of EF and 0.18 ± 0.01 d^-1^ in the presence of DF, while in the control condition growth rate corresponded to 0.39 ± 0.01 d^-1^) (Bonferroni’s test; *p* < 0.05). In contrast, the exposure to 20-fold concentrated vesicular fractions did not affect chlorophyte growth **(Figures 4A and 5A)**.

**Figure 4:**
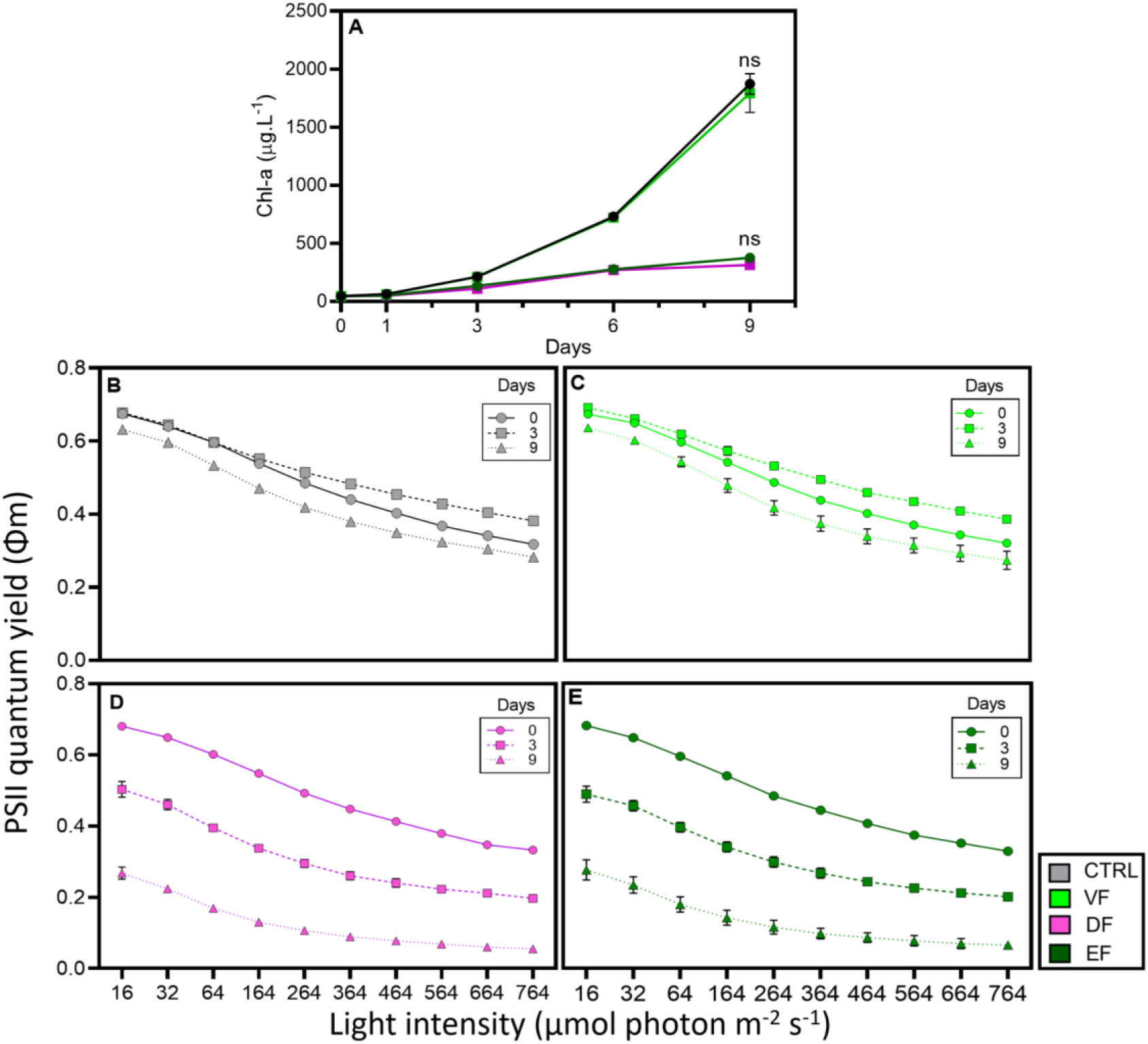
Allelopathic effects of exudate fractions of *R. raciborskii* T3 strain on *M. capricornutum* growth and photosynthetic activity. (A) Chlorophyll concentration curves of *M. capricornutum* (two-way RM ANOVA with Bonferroni *post hoc* test, where *ns* denotes no significant difference). (B-E) Maximum quantum yield of PSII derived from photosynthesis-irradiance curves of *M. capricornutum* cultures exposed to exudate fractions of the *R. raciborskii* T3 strain. (B) Control (ASM-1 + 1.43% PBS); (C) vesicular fractions (VF); (D) dissolved fractions (DF); (E) extracellular fractions (EF).

**Figure 5:**
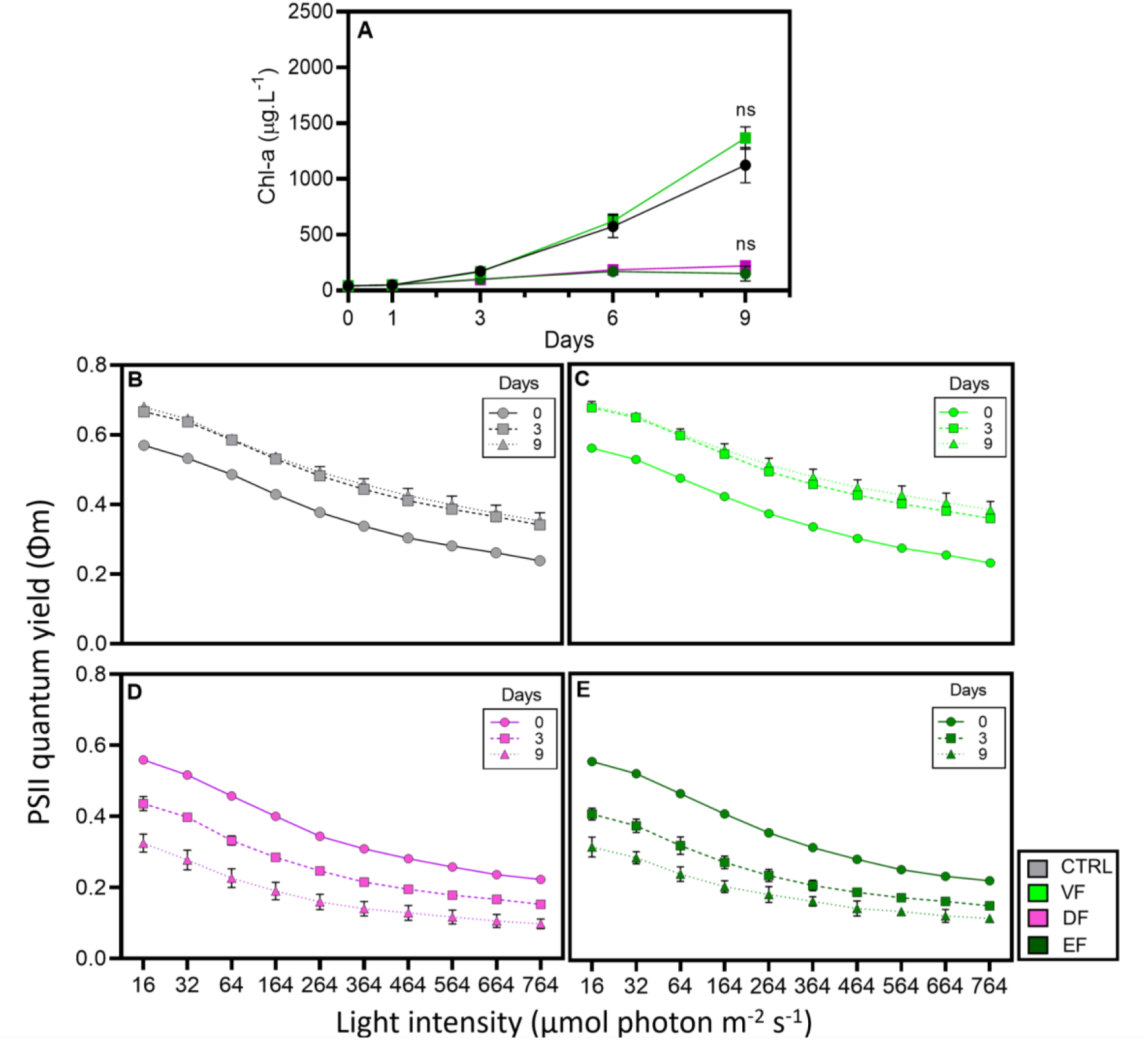
Allelopathic effects of exudate fractions of *R. raciborskii* CY-10 strain on *M. capricornutum growth* and photosynthetic activity. (A) Chlorophyll concentration curves of *M. capricornutum* (two-way RM ANOVA with Bonferroni *post hoc* test, where *ns* denotes no significant difference). (B-E) Maximum quantum yield of PSII derived from photosynthesis-irradiance curves of *M. capricornutum* cultures exposed to exudate fractions of the *R. raciborskii* CY-10 strain. (B) Control (ASM-1 + 1.43% PBS); (C) vesicular fractions (VF); (D) dissolved fractions (DF); (E) extracellular fractions (EF).

The maximum quantum yield of photosystem II (PSII) was represented by light-response curves **(Figures 4B-E and 5B-E)**. A higher stress to increased light pulses (at each 10s) and a remarkable reduction in PSII quantum yield over incubation time was observed in the chlorophyte cultures exposed to EF and DF obtained from both *R. raciborskii* strains, reinforcing the cyanobacterial exudate allelopathic effect. However, no reduction in quantum yield was detected over time in chlorophyte cultures exposed to VF, with curve profiles similar to control (ASM-1).

Further photosynthetic parameters such as light saturation (*Ik*), maximum electron transport rate (ETR_max_), and photosynthetic efficiency (α) were also determined in *M. capricornutum* cultures exposed to the exudate fractions of *R. raciborskii* **(Supplementary Tables S2 and S3)**. All these parameters were decreased following 9 days of exposure to allelochemicals. Again, the chlorophyte responded similarly when exposed to EF or DF (two-way RM-ANOVA, *p* > 0.05). The fraction containing concentrated extracellular vesicles (VF) did not alter the photosynthetic activity of *M. capricornutum* as compared to the control.

To evaluate if the allelopathic effect observed for EF and DF could be attributed to STX, *M. capricornutum* cultures were exposed to purified toxin preparations in final concentrations corresponding to those measured for the EF of each *R. raciborskii* strain (0.2 ng.mL^-1^ and 2 ng.mL^-1^ for T3 and CY-10, respectively). EFs obtained from each strain were used as positive controls for the effect of other allelochemicals different from saxitoxins **(Figure 6)**.

**Figure 6:**
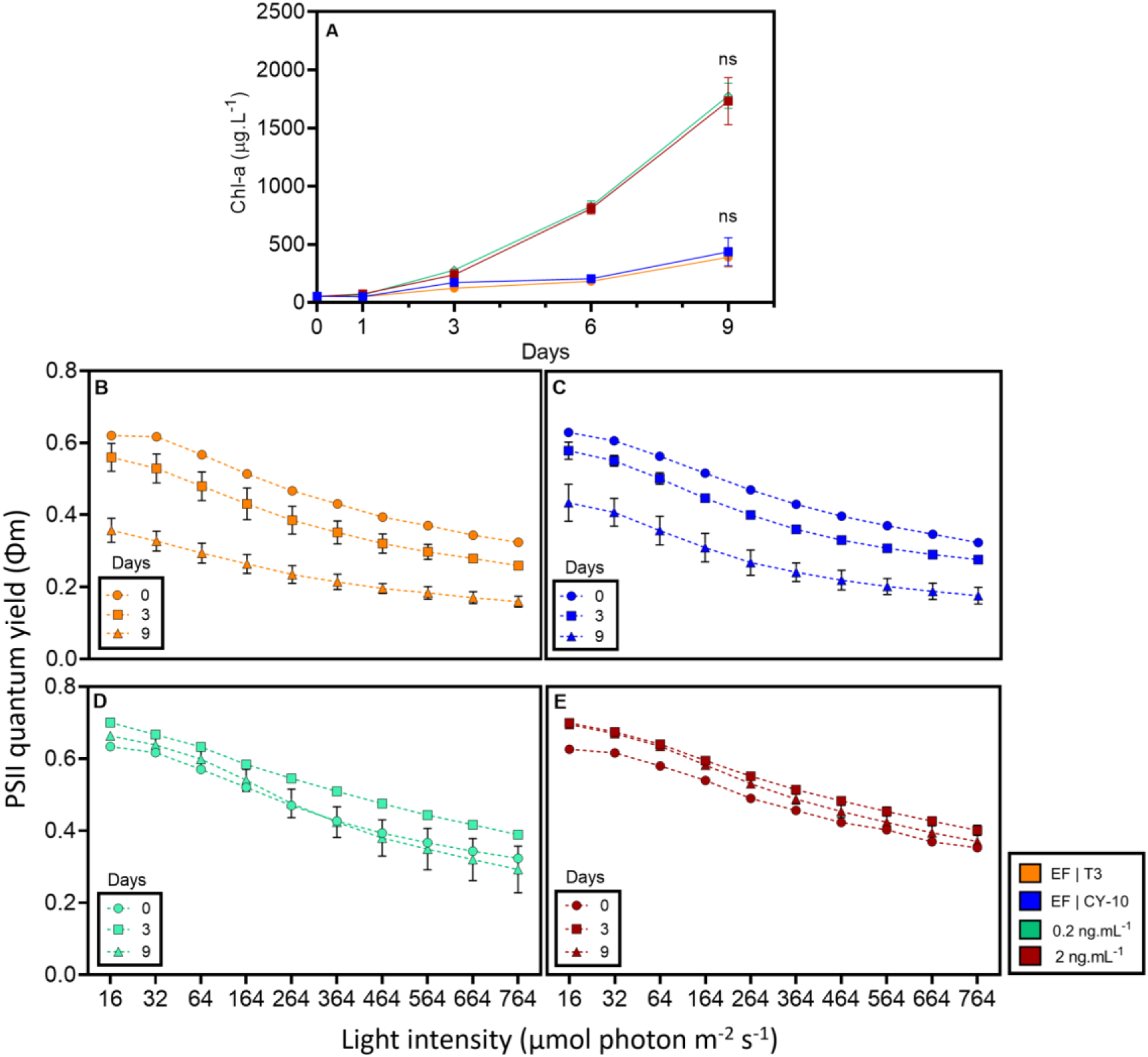
Effects of purified STX and exudate fractions of *R. raciborskii* on *M. capricornutum* growth and photosynthetic activity. (A) Chlorophyll concentration curves of *M. capricornutum* (two-way RM ANOVA with Bonferroni *post hoc* test, where *ns* denotes no significant difference). (B-E) Maximum quantum yield of PSII derived from the photosynthesis-irradiance curves of *M. capricornutum*. (B) Cultures exposed to extracellular fraction (EF) of the T3 strain ; (C) Cultures exposed to the extracellular fraction of the CY-10 strain; (D) Cultures exposed to 0.2 ng.mL^-1^ of STX; (E) Cultures exposed to 2.0 ng.mL^-1^ of STX.

The presence of STXs up to 2 ng.mL^-1^ did not affect algal growth as assessed by chlorophyll-α content **(Figure 6A)** or biomass yield **(Supplementary Table S4).** Growth rates in cultures treated with 2 ng.mL⁻¹ and 0.2 ng.mL⁻¹ STX were 0.39 ± 0.01 d^-1^ and 0.41 ± 0.01 d^-1^, respectively, similar to the control group (ASM-1), with a growth rate of 0.40 ± 0.02. In contrast, algal cultures exposed to EFs from T3 and CY-10 strains exhibited growth rates of .0.25 ± 0.03 d^-1^ and 0.26 ± 0.03 d^-1^, respectively,

The maximum quantum yield of photosystem II (PSII), represented by light-response curves, showed that STXs did not affect algal photosynthesis, while EF reduced PSII quantum yield **(Figure 6B-E)**. The other assessed photosynthetic parameters (α, ETR_max_, *Ik*) changed accordingly, confirming the inhibitory allelopathic effect of EFs and the null effect of STX **(Table S5)**.

## Discussion

In our findings, exudates of two saxitoxin-producing *Raphidiopsis raciborskii* strains were characterized for extracellular vesicle (EVs) presence by transmission electron microscopy (TEM) revealing particles with around 100 nm in size. The production of EVs by *R. raciborskii* was first documented by Zarantonello et al. (2018) which by using TEM revealed structures with 20 to 300 nm (mostly 140-300 nm) in diameter that originated from the outer membrane. Here, besides TEM, Nanoparticle Tracking Analysis (NTA) evidenced EVs ranging from 80 to 350 nm (median 204 nm). Thus, although a variation in EV size is expected for different cyanobacterial strains (Biller et al., 2023), a similar size distribution was revealed for *R. raciborskii* considering the two strains in the present study (T3 and CY-10) and that previously characterized (CYRF-01) by Zarantonello et al. (2018). Similarly, for other filamentous cyanobacteria, the evaluation of EV sizes by TEM indicated mostly structures ∼100-200 nm for species with diverse cell volumes such as the freshwater planktonic *Anabaena* sp. PCC 7120 (Oliveira et al., 2016), the freshwater *Leptolyngbya boryana* (Usui et al., 2022; 2024), the marine *Jaaginema litorale* (Brito et al., 2017) and the symbiont *Anabaena azollae*, during differentiation into akinetes (Zheng et al., 2009). Considering the low number of cells analyzed by TEM in these studies, it is expected that future investigations using NTA results will better represent EV size distributions, reflecting the diversity of vesicles produced by populations of cells, as demonstrated for the picocyanobacteria *Synechococcus* and *Prochlorococcus*, whose EV diameters range of 50-100 nm (Biller et al., 2014, 2023). This will clarify if there is a proportional relationship between cell biovolume and EV biovolume.

On average, the T3 strain released approximately twice as many EVs per cell as the CY-10 strain, suggesting that EV production may vary between strains. Similar findings have also been observed for the marine cyanobacterium *Prochlorococcus* (Biller et al., 2023). These values are comparable to those obtained for the *R. raciborskii* CYRF-01 strain (Zarantonello et al., 2018), although in that study a different measure was taken, by estimating the number of particles per cell section based on TEM images. This resulted in 4-6 EVs per cell section, indicating that tens of EVs would be produced per cell.

In the present study, we used two STX-producing *R. raciborskii* strains to test whether EVs could participate in toxin release. Although STXs are predominantly intracellular in cyanobacteria, they can be detected in the extracellular environment, albeit at volumetric concentrations about 10 times lower than those detected in the cellular fraction (Vilar et al., 2021). The analysis of *sxt* gene cluster components in *Raphidiopsis* genomes indicated that the toxin can be exported. The *sxtM* and *sxtF* genes encode transporter proteins from the MATE (multidrug and toxic compound extrusion) family (Soto-Liebe et al., 2013). Predicted protein structures of SxtF and SxtM from *R. brookii* D9 were elucidated and a binding domain for STX analogues was proposed in the protein cytoplasmic portion (Soto-Liebe et al., 2013). Possibly, STXs could be transported to the periplasm and then secreted into EVs. For both strains, the toxin concentrations in the extracellular fraction (EF, total exudate containing vesicles) was similar to that found in the dissolved fraction (DF, exudate after removal of the vesicles by filtration), suggesting that, in the extracellular environment the toxin is mainly found in the free, dissolved form. Nonetheless, STXs were also detected in the vesicular fraction (VF), albeit at concentrations much lower than those recorded in the other fractions.

Considering that the EV volume (∼0.004 μm^3^) is much smaller than the average cell volume (∼45 μm^3^), toxin concentrations were converted to quota in relation to biovolume. This resulted in a quota per vesicle approximately 50 times lower than the cellular quota for T3 and 17 times lower for CY-10. The distribution of STX analogues in the extracellular fractions differed from that found in the intracellular fraction. While in cells the predominant analogue was neoSTX, in the extracellular fraction STX showed greater relative abundance. Although the reasons for this are not clear, it is likely that some interconversion of analogues may occur in the extracellular environment, e.g., through chemical alteration or enzymatic activity (Begoña Ben-Gigirey; Adriano Villar-González, 2008).

It must be considered that the quantification of toxins in EVs presents limitations. One point is the possible loss of toxins from the EV to the medium during the EV isolation process, given the manipulation involved during EV isolation and the hydrophilic characteristic of the STX molecule (Wiese et al., 2010). Furthermore, since the cultures used to obtain vesicles are not in axenic conditions, EV preparations could include a fraction of EVs from the microbiota associated with the cyanobacteria, although the latter largely predominated in terms of biomass. These interfering factors may have led to an underestimation of STX concentrations in VFs. Therefore, even at low concentrations, we conclude that EVs contain saxitoxins and that this can constitute a secretion path for toxins and other secondary metabolites, as demonstrated in other bacteria (Schwechheimer & Kuehn, 2015; Toyofuku et al., 2019).

The *R. raciborskii* T3 strain toxin profile characterized in the present study was in accordance with that described previously (D’Agostino et al., 2019), with the predominance of the neoSTX analogue. Total STX concentration was twelve-fold higher in the intracellular fraction than in the extracellular. For the CY-10 strain, the toxin profile was first described in the present study, comprising mainly neoSXT and a minor contribution of STX, and GTX 1 and 2. Total intracellular STX concentration was five-fold higher than the extracellular concentration. *R. raciborskii* CY-10 presented a higher cellular quota of STXs, which suggests a greater relative toxicity compared to T3.

Given that the EVs could contain secondary metabolites, we hypothesized that they could participate in the delivery of allelochemicals. This idea was not confirmed, since the addition of purified, concentrated EV preparations to cultures of *Monoraphidium capricornutum* did not inhibit growth or photosynthesis. In similar assays, both the extracellular fraction (with vesicles) and the filtrate (without vesicles) exerted a similar inhibitory effect on the growth and photosynthetic parameters of the chlorophyte. Previous studies described the allelopathic effect of *R. raciborskii* exudates on phytoplankton components and allelopathy is considered a relevant contributor to the expansion of this species in freshwater environments (Antunes et al., 2015; Figueredo et al., 2007). Culture filtrates (exudates) from several *R. raciborskii* strains (for which toxin production was not characterized) inhibited the photosynthetic activity of the green algae *Coelastrum sphaericum,* and the exudate from a single *R. raciborskii* strain inhibited photosynthesis of diverse species, such as the cyanobacteria *Microcystis aeruginosa* and *Microcystis wesenbergii*, and the chlorophyte *Monoraphidium contortum* (Figueredo et al., 2007). The exudate of *R. raciborskii* strains inhibited the growth of *Ankistrodesmus falcatus* (Antunes et al., 2012; Leão et al., 2009a) and *M. aeruginosa* (Rzymski et al., 2014). In another study, exudates obtained from mixed cultures of *R. raciborskii* and *M. aeruginosa* inhibited the growth of *M. aeruginosa* and induced colony formation, suggesting that competition may trigger the allelopathic effect (Mello et al., 2012).

The chemical composition of *R. raciborskii* exudates is still uncharacterized, mainly regarding their allelopathic inhibitory effect. However, our study moves forward by fractionating the cyanobacterial exudates and testing whether an EVs enriched fraction or a dissolved fraction contributes to the already documented allelopathic effect of the cyanobacterium on other phytoplankton. In the present study, the inhibitory effect of the exudates obtained from the two toxic strains was preserved in the dissolved fraction (<100 kDa). This led to the question of whether the toxin present in the dissolved fraction could exert an allelopathic effect. In this line, further tests with purified STX in concentrations similar to those found in the extracellular medium to *M. capricornutum* cultures did not affect growth or photosynthesis. Similar results have been reported using other target species. Addition of purified STX (2nM up to 128nM or 0.6-40 μg L^−1^) did not affect the physiology of the green algae *Chlamydomonas reinhardtii*. The toxin did not inhibit cell division or PSII activity, and did not alter ROS levels or cell viability (Perreault et al., 2011). In a subsequent study, among diverse physiological aspects assessed, only a negative effect on the activity of antioxidant enzymes of *C. reinhardtii* was observed upon exposure to pure STX (3.0 nM) (Melegari et al., 2012). Using a different approach, it was demonstrated that *R. raciborskii* crude cell extracts of both STX-producing and non-producing strains inhibited the growth of the cyanobacteria *M. aeruginosa* and *M. wesenbergii*, and of the green algae *Tetradesmus lagerheimii* (formerly *Scenedesmus acuminatus*) (Bittencourt-Oliveira et al., 2016). A more pronounced effect was noted in the case of STX+ extracts, suggesting that STX was related to toxic effects on phytoplankton. However, the effects observed in STX-extracts suggested that other metabolites may act as allelochemicals.

In the present study, it was reinforced that the exudates of *R. raciborskii* (containing molecules <100 kDa) have an allelopathic effect on *M. capricornutum* by inhibiting photosynthesis and growth. A reduction in photosynthetic activity was apparent from the decrease in values of PSII quantum yield, the light saturation point of photosynthesis (*Ik*), maximum electron transport rate (ETR_max_), and photosynthetic efficiency (α). These results are similar to those of previous investigations that tested allelopathic interactions using *Monoraphidium* as a target organism. Cell-free lake water or exudates of *R. raciborskii* in culture inhibited photosynthesis of *Monoraphidium contortum*, decreasing maximal ETR values (Figueredo et al., 2007). Filtrates from *Synechococcus* sp. inhibited the synthesis of photosynthetic pigments (chlorophyll-α and carotenoids) by *Monoraphidium* sp. and other green algae (Konarzewska et al., 2020). Indeed, for interactions between photoautotrophs in aquatic habitats, allelochemicals are mainly inhibitors of photosynthesis, reducing the electron transport rate and quantum yield of photosystem II, which is submitted to oxidative damage (Gross, 2003; Pei et al., 2020).

In summary, extracellular vesicles present in the exudate of toxic strains of the cyanobacterium *Raphidiopsis raciborskii* were characterized. Saxitoxins were first detected in cyanobacterial EVs. A possible biological function of EVs in this species of cyanobacteria have been suggested as part of the response to biotic stress, based on the observation of increased vesiculation in *R. raciborskii* upon interaction with *Microcystis aeruginosa* (Zarantonello et al., 2018), which could be related to allelopathic interactions. Here, the role of EVs in the allelopathic interaction with another phytoplankton species was tested, using the microalgae *Monoraphidium capricornutum* as a model, but no effect was observed. Therefore, the function of the release of these structures by *R. raciborskii* remains unknown and deserves further investigation.

## Acknowledgments

The authors would like to thank Dr Carolina Keim for the support for obtaining the Transmission Electron Microscopy images and also the technical assistance of the National Center for Structural Biology and Bioimaging, Federal University of Rio de Janeiro. We would also like to thank Dr Susana Frases for her valuable advice regarding nanoparticle analysis. The study was carried out with the financial support of the Conselho Nacional de Desenvolvimento Científico e Tecnológico – CNPq (Brazil) (award number: 408525/2023-1), Fundação Carlos Chagas Filho de Amparo à Pesquisa do Estado do Rio de Janeiro (FAPERJ, Brazil) and the Coordenação de Aperfeiçoamento de Pessoal de Nível Superior (CAPES, Brazil) for C.P.N. MD scholarship.

