## Supplemental material for "Extracellular vesicles from saxitoxin-producing strains of the cyanobacterium *Raphidiopsis raciborskii* and investigation of their potential allelopathic effect"

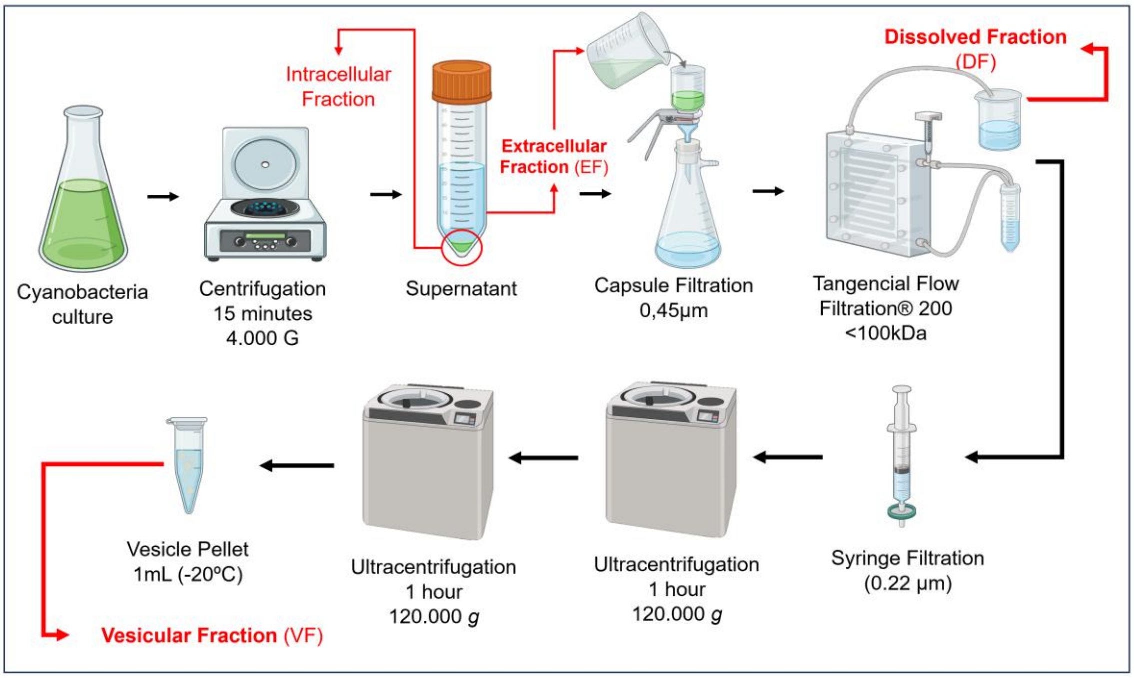


**Supplementary Figure 1:** Scheme of the methodology used to obtain EVs from *Raphidiopsis raciborskii* cultures. Methodology adapted from Biller (2022)

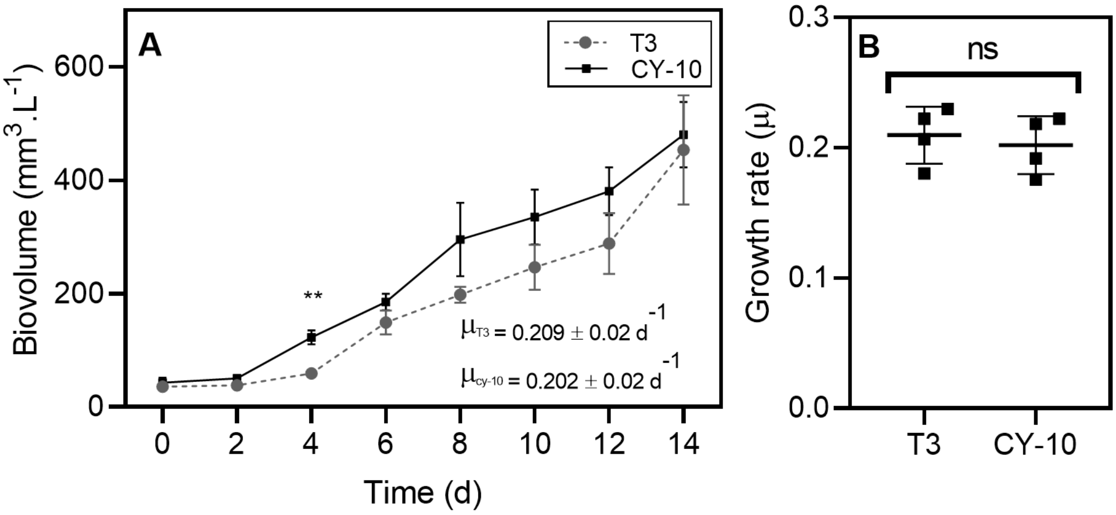


**Supplementary Figure 2:** Growth of *R. raciborskii* strains. (A) Growth curves were established for T3 (dashed line) and CY-10 (solid line) strains over 14 days. An increase in biomass initiated between the 2nd and 4th day of incubation and continued onward. From the growth curves, the exponential growth phase was determined to obtain extracellular vesicle (EV) preparations, ensuring a higher biomass of viable cells and, consequently, a high vesicle yield for subsequent analyses. (B) Specific growth rates of T3 and CY-10 strains. No significant differences were observed between the strains (ns = not significant, Student's t-test, $p > 0.05$) ($N=4$).

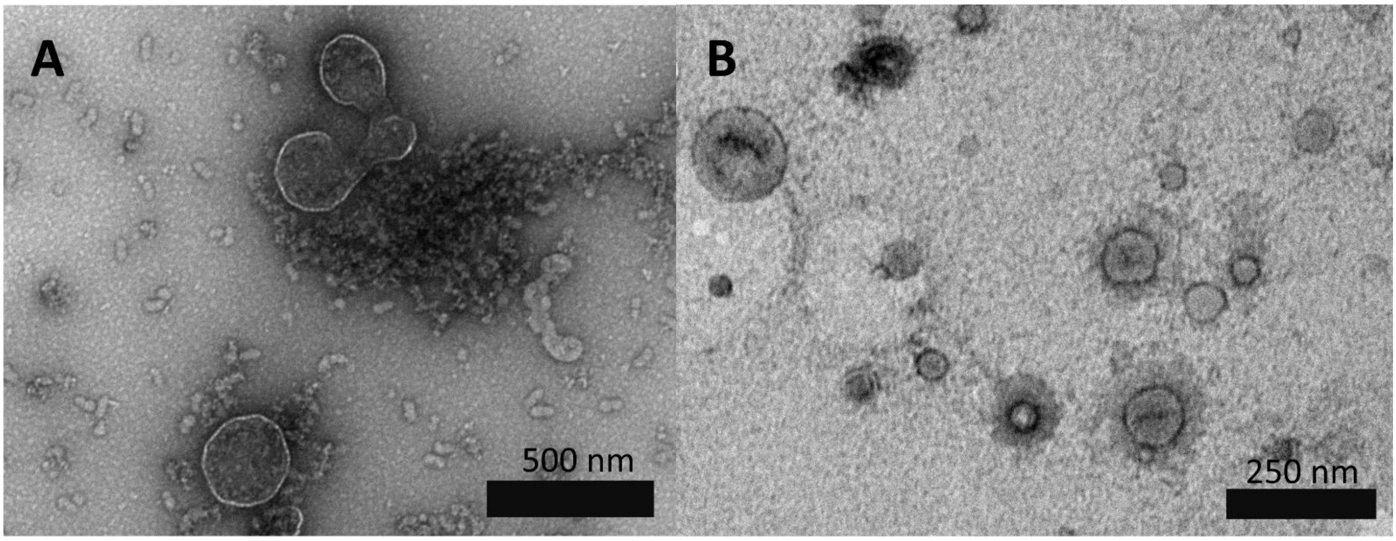


**Supplementary Figure 3**: TEM images using negative stain of vesicle preparations obtained from *R. raciborskii* strains, (A) T3 and (B) CY-10.

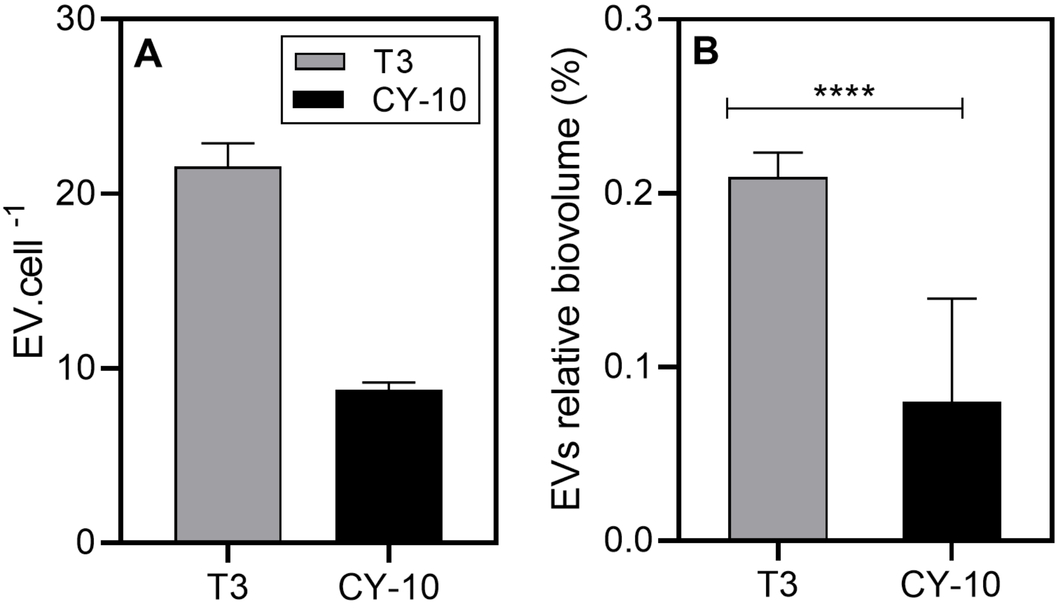


**Supplementary Figure 6:** EV production per cell for the T3 and CY-10 strains. (A) Number of vesicles per number of cells (B) Biovolume of vesicles per biovolume of cells (**** significant difference, Student's t-test, $p < 0.0001$)

**Table S1**: Growth of *Monoraphidium capricornutum* exposed to exudate fractions of *R. raciborskii* strains. Growth yield is defined as the number of cells at the final time (9 days) divided by the number of cells at the initial time, while biomass yield corresponds to the number of cells at the final time (9 days) minus the initial number of cells. Comparisons among the fractions were performed for each strain and different letters indicate significant differences (Bonferroni's test, $p < 0.05$; Mean ± SD).

| Strain | Treatment | Growth yield (fold-change) | Cells yield (cells.mL$^{-1}$) |
|---|---|---|---|
| T3 | Control (ASM-1) | 47.2 (± 2.83)$^{A}$ | $4.19\times10^{6}$ (± $2.57\times10^{5}$)$^{A}$ |
| | EV | 46.9 (± 7.60)$^{A}$ | $3.90\times10^{6}$ (± $6.45\times10^{5}$)$^{A}$ |
| | EF | 8.8 (± 1.23)$^{B}$ | $6.84\times10^{5}$ (± $1.07\times10^{5}$)$^{B}$ |
| | DF | 6.4 (± 0.15)$^{B}$ | $5.26\times10^{5}$ (± $1.45\times10^{4}$)$^{B}$ |
| CY-10 | Control (ASM-1) | 93.6 (± 5.58)$^{A}$ | $3.64\times10^{6}$ (± $2.20\times10^{5}$)$^{A}$ |
| | EV | 96.8 (± 9.61)$^{A}$ | $4.47\times10^{6}$ (± $4.48\times10^{5}$)$^{B}$ |
| | EF | 4.5 (± 0.48)$^{B}$ | $1.60\times10^{5}$ (± $2.24\times10^{4}$)$^{C}$ |
| | DF | 6.8 (± 1.24)$^{B}$ | $2.36\times10^{5}$ (± $5.06\times10^{4}$)$^{C}$ |

**Table S2**: Photosynthetic parameters of *Monoraphidium capricornutum* exposed to extracellular fractions of *R. raciborskii* T3 strain. Yield – relative quantum yield of PSII during a saturation pulse of 16 PAR (photosynthetically active radiation); $ETR_{max}$ – maximum electron transport rate; Ik – light saturation parameter; and Alpha (α) – light-harvesting efficiency. Comparisons among the fractions were performed and different letters indicate significant differences (Bonferroni's test, $p < 0.05$; Mean ± SD).

| Strain | Time | Fraction | Yield (*Fv'/Fm'*) | Alpha (µmol fóton.m$^{-2}$ s$^{-1}$) | $ETR_{max}$ (µmol e.m$^{-2}$ s$^{-1}$) | Ik (µmol photon.m$^{-2}$ s$^{-1}$) |
|---|---|---|---|---|---|---|
| T3 | Day 0 | CTRL | 0.68 (± 0)$^{A}$ | 0.28 (± 0.004)$^{A}$ | 188 (± 4.80)$^{A}$ | 671.55 (± 26.80)$^{A}$ |
| | | VF | 0.69 (± 0)$^{A}$ | 0.284 (± 0.000)$^{A}$ | 189.75 (± 2.90)$^{A}$ | 667.9 (± 11.59)$^{A}$ |
| | | EF | 0.68 (± 0)$^{A}$ | 0.284 (± 0.002)$^{A}$ | 196.3 (± 4.24)$^{A}$ | 691.95 (± 22.41)$^{A}$ |
| | | DF | 0.68 (± 0)$^{A}$ | 0.285 (± 0.002)$^{A}$ | 200.3 (± 0.42)$^{A}$ | 703.05 (± 4.31)$^{A}$ |
| | Day 9 | CTRL | 0.64 (± 0)$^{B}$ | 0.261 (± 0.000)$^{B}$ | 153.25 (± 6.54)$^{B}$ | 586.7 (± 24.45)$^{B}$ |
| | | VF | 0.6425 (± 0.009)$^{B}$ | 0.266 (± 0.002)$^{B}$ | 143.25 (± 19.83)$^{B}$ | 538.125 (± 73.27)$^{B}$ |
| | | EF | 0.31 (± 0.008)$^{C}$ | 0.124 (± 0.002)$^{C}$ | 26.1 (± 1.88)$^{C}$ | 209.85 (± 10.89)$^{C}$ |
| | | DF | 0.255 (± 0.031)$^{D}$ | 0.100 (± 0.010)$^{C}$ | 15.875 (± 2.77)$^{D}$ | 153.05 (± 25.95)$^{C}$ |

**Table S3**: Photosynthetic parameters of *Monoraphidium capricornutum* exposed to extracellular fractions of *R. raciborskii* CY-10 strain. Yield – relative quantum yield of PSII during a saturation pulse of 16 PAR (photosynthetically active radiation); $ETR_{max}$ – maximum electron transport rate; Ik – light saturation parameter; and Alpha (α) – light-harvesting efficiency. Different letters indicate significant diferences (Bonferroni's test, $p < 0.05$; Mean ± SD).

| Strain | Time | Fraction | Yield (*Fv'/Fm'*) | Alpha (μmol fóton.m$^{-2}$ s$^{-1}$) | ETRmax (μmol e.m$^{-2}$ s$^{-1}$) | Ik (μmol photon.m$^{-2}$ s$^{-1}$) |
|---|---|---|---|---|---|---|
| CY-10 | Day 0 | CTRL | 0.58 (± 0)$^{A}$ | 0.237 (± 0.001)$^{A}$ | 124.6 (± 1.69)$^{A}$ | 524.5 (± 4.38)$^{A}$ |
| | | VF | 0.57 (± 0)$^{A}$ | 0.234 (± 0.002)$^{A}$ | 124.8 (± 4.95)$^{A}$ | 532.35 (± 26.09)$^{A}$ |
| | | EF | 0.57 (± 0)$^{A}$ | 0.234 (± 0.004)$^{A}$ | 110.45 (± 4.59)$^{A}$ | 472.25 (± 11.38)$^{A}$ |
| | | DF | 0.57 (± 0)$^{A}$ | 0.231 (± 0.001)$^{A}$ | 108.75 (± 3.89)$^{A}$ | 468.85 (± 15.48)$^{A}$ |
| | Day 9 | CTRL | 0.69 (± 0.008)$^{B}$ | 0.279 (± 0.003)$^{B}$ | 230.375 (± 29.95)$^{B}$ | 825 (± 105.58)$^{B}$ |
| | | VF | 0.68 (± 0.005)$^{B}$ | 0.280 (± 0.003)$^{B}$ | 273.1 (± 37.42)$^{B}$ | 975.25 (± 127.44)$^{B}$ |
| | | EF | 0.32 (± 0.026)$^{C}$ | 0.128 (± 0.009)$^{C}$ | 53.475 (± 7.93)$^{C}$ | 414.65 (± 44.10)$^{C}$ |
| | | DF | 0.33 (± 0.025)$^{C}$ | 0.1312 (± 0.010)$^{C}$ | 38.975 (± 8.29)$^{C}$ | 294.375 (± 40.68)$^{D}$ |

**Table S4**: Growth of *Monoraphidium capricornutum* exposed to saxitoxins. Relative growth yield is defined as the number of cells at the final time (9 days) divided by the number of cells at the initial time, while biomass yield corresponds to the number of cells at the final time (9 days) minus the initial number of cells. Comparisons among the fractions were performed and different letters indicate significant differences (Bonferroni's test, $p < 0.05$; Mean ± SD).

| Treatment | Growth yield (fold-change) | Cells yield (cells.mL$^{-1}$) |
|---|---|---|
| EF\|CY-10 | 9.5 (± 1.66)$^{A}$ | $1.02\times10^{6}$ (± $2.00\times10^{5}$)$^{A}$ |
| EF\|T3 | 7.3 (± 1.80)$^{A}$ | $8.16\times10^{5}$ (± $2.32\times10^{5}$)$^{A}$ |
| STX\|2.0 ng.ml$^{-1}$ | 46.7 (± 5.46)$^{B}$ | $5.87\times10^{6}$ (± $7.01\times10^{5}$)$^{B}$ |
| STX\|0.2 ng.ml$^{-1}$ | 40.5 (± 11.04)$^{B}$ | $5.39\times10^{6}$ (± $1.51\times10^{6}$)$^{B}$ |

**Table S5**: Photosynthetic parameters of *Monoraphidium capricornutum* exposed to the extracellular fractions of *R. raciborskii* strains or purified saxitoxins. Yield – relative quantum yield of PSII during a saturation pulse of 16 PAR (photosynthetically active radiation); $ETR_{max}$ – maximum electron transport rate; Ik – light saturation parameter; and Alpha (α) – light-harvesting efficiency. Different letters indicate significant diferences (Bonferroni's test, $p < 0.05$; Mean ± SD).

| Time | Fraction | Yield (*Fv'/Fm'*) | Alpha (µmol fóton.$m^{-2}$ $s^{-1}$) | ETRmax (µmol e.$m^{-2}$ $s^{-1}$) | Ik (µmol photon.$m^{-2}$ $s^{-1}$) |
|---|---|---|---|---|---|
| Day 0 | EF \| CY-10 | 0.63 (± 0.017)$^{A}$ | 0.259 (± 0.006)$^{A}$ | 205.00 (± 5.65)$^{A}$ | 807.45 (± 23.83)$^{A}$ |
| | EF \| T3 | 0.64 (± 0.01)$^{A}$ | 0.268 (± 0.004)$^{A}$ | 213.3 (± 14.28)$^{A}$ | 789.15 (± 32.88)$^{A}$ |
| | STX \| 2.0 ng.$mL^{-1}$ | 0.65 (± 0.005)$^{A}$ | 0.254 (± 0.02)$^{A}$ | 236.10 (± 15.84)$^{A}$ | 885.95 (± 57.91)$^{A}$ |
| | STX \| 0.2 ng.$mL^{-1}$ | 0.64 (± 0.01)$^{A}$ | 0.268 (± 0.00)$^{A}$ | 206.50 (± 11.03)$^{A}$ | 775.75 (± 50.98)$^{A}$ |
| Day 9 | EF \| CY-10 | 0.45 (± 0.04)$^{B}$ | 0.180 (± 0.02)$^{B}$ | 86.1 (± 11.87)$^{B}$ | 476.6 (± 14.61)$^{B}$ |
| | EF \| T3 | 0.33 (± 0.04)$^{B}$ | 0.143 (± 0.01)$^{B}$ | 95.33 (± 14.06)$^{B}$ | 637.03 (± 32.17)$^{B}$ |
| | STX \| 2.0 ng.$mL^{-1}$ | 0.70 (± 0.005)$^{C}$ | 0.290 (± 0.0)$^{C}$ | 243.73 (± 31.15)$^{C}$ | 839.97 (± 114.24)$^{BC}$ |
| | STX \| 0.2 ng.$mL^{-1}$ | 0.67 (± 0.02)B$^{C}$ | 0.280 (± 0.0)$^{C}$ | 187.43 (± 14.7)$^{C}$ | 679.8 (± 34.27)$^{BC}$ |
